# Quantifying the phenotypic information in mRNA abundance

**DOI:** 10.1101/2022.02.23.481668

**Authors:** Evan Maltz, Roy Wollman

## Abstract

Quantifying the dependency between mRNA abundance and downstream cellular phenotypes is a fundamental open problem in biology. Advances in multimodal single cell measurement technologies provide an opportunity to apply new computational frameworks to dissect the contribution of individual genes and gene combinations to a given phenotype. Using an information theory approach, we analyzed multimodal data of the expression of 83 genes in the Ca^2+^ signaling network and the dynamic Ca^2+^ response in the same cell. We found that the overall expression levels of these 83 genes explain approximately 60% of Ca^2+^ signal entropy. The average contribution of each single gene was 16%, revealing a large degree of redundancy between genes. Using different heuristics we estimated the dependency between the size of a gene set and its information content, revealing that on average a set of 53 genes contains 90% of the information about Ca^2+^ signaling within the cellular transcriptional state. Our results provide the first direct quantification of information content about complex cellular phenotype that exists in mRNA abundance measurements.

## Introduction

Cellular phenotypes emerge from many regulated interactions between various components. Rates of synthesis and degradation determine the instantaneous abundances of different biological molecules. These kinetic rates are themselves a property of regulatory interactions between biomolecules creating multilayered feedback networks (El-Samad, 2021). Both the dynamic and instantaneous abundances of biomolecules are key determinants of cellular phenotypes, underlying their ability to make different decisions given the same stimulus (Purvis & Lahav, 2013; Perkins & Swain, 2009; Cheong *et al*, 2011). The ability to systematically measure the abundance of large sets of different biomolecules such as mRNA and proteins enables the determination of regulatory strengths across different nodes of these complex networks. Pioneering work in *E. coli* based on instantaneous single cell measurements of mRNA and protein copy numbers reveal a surprisingly low correlation coefficient of r=0.01 ± 0.03 across 129 highly expressed genes (Taniguchi *et al*, 2010). The lack of correlation between mRNA and protein in *E. coli* might be due to their small size and magnitude of temporal fluctuation in mRNA levels. However, more recent advances in multimodal assays in mammalian cells also identified low correlations between the abundances of most proteins and corresponding mRNAs (Stoeckius *et al*, 2017; Gong *et al*, 2017; Schulz *et al*, 2018; Mair *et al*, 2020; Darmanis *et al*, 2016). This low correlation appears to contradict intuition that protein and mRNA levels should strongly correspond within cells because of the dependency suggested by the central dogma (Liu *et al*, 2016). An alternative hypothesis is that the majority of regulatory steps and phenotypically-relevant information lie in post transcriptional processes. Post-transcriptional regulation can modulate both protein activity and abundance via protein interactions, post-translational modifications, RNA interactions/structure, and more. Stochastic processes also obscure the importance of molecular composition to phenotypic outcomes (Balázsi *et al*, 2011; Perkins & Swain, 2009; Cheong *et al*, 2011). Yet, many studies have pointed to differences in mRNA levels among clonal cells to explain differences in cellular phenotypes (Shaffer *et al*, 2020; Emert *et al*, 2021). These observations highlight a need for a better framework to address fundamental questions: Does mRNA abundance matter? What fraction of the information about cellular phenotype is determined by mRNA abundance, and what fraction is due to post-transcriptional regulation?

Quantifying the information content in mRNA abundance about cellular phenotypes is technically and computationally challenging due to the many layers of complex interactions in cellular networks (Macaulay *et al*, 2017). Phenotypic information displayed in clonal cells is controlled by molecular composition, stochastic factors, intermediate regulation, and crosstalk (Azeloglu & Iyengar, 2015). Many approaches have been developed to disentangle these complex and distributed dependencies. Feature engineering has been one powerful tool to reveal interpretable characteristics of signaling dynamics, finding multiple motifs that encode information about stimulus dose and type (Adelaja *et al*, 2021; Zhang *et al*, 2017; Wong *et al*, 2019; Hafner *et al*, 2017; Nelson *et al*, 2004). However, these features do not capture all the information in complex dynamics, which are difficult to study and fully recapitulate in mechanistic models (Myers *et al*, 2021). Another common approach is to perform dimensionality reduction and/or clustering to integrate different modalities (Subramanian *et al*, 2020; Kinnunen *et al*, 2021). Several studies have clustered groups of genes or cells based on signal patterns to reveal general mechanisms of how signaling dynamics affect transcription (Lane *et al*, 2017; Hafner *et al*, 2017). However, it is still unknown how differences in arbitrary sets of transcripts relate to dynamic signals in the same cells. Signaling phenomena may emerge due to differences among many combinations of genes, which may be missed when simplified to either individual genes or gene clusters. Single-cell network states are notoriously difficult to fully measure, and insights about the relationships between many components requires high dimensional and multimodal data from the same cells (Adelaja *et al*, 2021; Spencer *et al*, 2009; Azeloglu & Iyengar, 2015; Macaulay *et al*, 2017). Although useful in many contexts, feature engineering, clustering, and dimensionality reduction are not guaranteed to capture all useful information.

Directly quantifying the relationship between many transcripts and a phenotype via an information theoretic approach can provide a direct measure of the importance of mRNA abundance. However, three challenges prevent the general use of information theory in quantifying information content in RNA abundance. 1) Biological feedbacks entangle mRNA abundance and cellular phenotype. Cellular phenotypes that emerge over long timescale, e.g. cellular differentiations, have a longer timescale than the lifetime of mRNA molecules that potentially determine the emerging phenotypes. In these cases, mRNA abundances themselves change dynamically adding additional complexities. 2) Quantification of importance of mRNA abundance requires paired measurements of mRNA and the emerging cellular phenotype in question, measurements that are technically challenging due to the destructive nature of mRNA quantification methods. 3) The statistical measures needed to answer these questions, entropy and mutual information, are notoriously hard to infer. Below, we discuss how these challenges could be addressed to provide direct quantification of the information content in mRNA abundance.

Ca^2+^ signaling is a useful model system to quantify the dependency between mRNA abundance and emerging cellular phenotypes. Ca^2+^ signaling is a system in which the emerging phenotype is faster than changes in mRNA abundance, allowing for timescale separation such that we can assume mRNA abundances are at a quasi-steady state. The dynamics of the Ca^2+^ signaling response to ATP is a well studied model system for environmental sensing, featuring one of the most ubiquitous and multifunctional pathways across cell types. An important role of Ca^2+^ signaling is the coordination of responses to changes in extracellular environment. In the physiological context of tissues, cell lysis causes an unusual local increase in extracellular ATP, among other molecules. This type of damage sensing relies on the purinergic cell surface receptors, P1 and P2, which detect adenosine and ATP respectively (Alves *et al*, 2018). The P2Y GPCR triggers a downstream signaling cascade via protein interactions. G_q_-GTP is released from the P2Y receptor where it can then bind to and activate phospholipase C (PLCβ). PLCβ cleaves PIP_2_ into IP_3_ and DAG, which facilitate signaling by binding to their respective receptors, IP3R and DAGR. The IP3R is embedded in the membrane of the endoplasmic reticulum and functions as a gated Ca^2+^ channel that releases Ca^2+^ into the cytoplasm upon IP_3_ binding. Cytoplasmic Ca^2+^ concentrations are kept relatively low at 50-100nM and spike up to 1uM during signaling with significant and rapid fluctuations producing unique dynamics in every cell (Bagur & Hajnóczky, 2017). Changes in cytoplasmic Ca^2+^ concentration over time (i.e. its signaling dynamics) have many emergent features like oscillations caused by coupling between positive and negative feedback loops (Azeloglu & Iyengar, 2015). Studies have shown these dynamics specifically propagate relevant environmental and stimulus information (Selimkhanov *et al*, 2014). While in the cytoplasm, Ca^2+^ regulates many signaling molecules, e.g. kinases and phophatases, through direct binding to Ca^2+^ binding domains such as the EF-hand and through binding to calmodulin isoforms that enables it to activate kinases like protein kinase C. These kinases affect many downstream transcriptional and protein-mediated responses that ultimately regulate cell behavior. The timescale of Ca^2+^ dynamics is significantly faster than the timescale of gene expression differentiation, allowing us to interpret a symmetric measure of dependency, such as mutual information, in a directed manner (Putney, 2012). Overall, the Ca^2+^ signaling pathway is a complex network with regulation at transcriptional and post-transcriptional levels, providing us with a great model system to dissect the phenotypic information content in mRNA abundances.

Precise measurements of dynamic single-cell, multimodal data have been collected to address these questions. Studies featuring multiomic image-based measurements have mostly focused on fixed cell measurements like immunofluorescence, spatial arrangement of cells in tissues, and chromatin structure (Wang *et al*, 2018; Zhang *et al*, 2021; Liu *et al*, 2021). Studies that have involved dynamic phenotypes were limited by the low sensitivity of scRNAseq (Lane et al. 2017) or had to focus on only a handful of genes (Lee et al. 2014). Nonetheless, methods are being developed to integrate live cell dynamics and reliable RNA quantification of hundreds of genes (Foreman & Wollman, 2020; Genshaft *et al*, 2021). Measuring the transcriptional state of the Ca^2+^ signaling pathway requires the quantification of the abundance of hundreds of genes. Multiplexed Error-Robust Fluorescence In Situ Hybridization (MERFISH) has been developed as a high-throughput single-cell method for accurately counting large numbers of transcripts (Moffitt *et al*, 2016). Because it is performed in situ, MERFISH can be combined with other imaging methods to create high dimensional, multimodal data consisting of both dynamic and instantaneous measurements. Combining transcriptomic and live-cell data offers unique insights into the role of dynamic regulation and sources of phenotypic information. The challenge of collecting high-dimensional, single cell, paired transcriptomic and signaling dynamics data has been successfully addressed in recent work (Foreman & Wollman, 2020). There, we demonstrated a single cell method for collecting paired measurements of live Ca^2+^ signaling dynamics and relevant gene expression using MERFISH. In that work, non-transformed epithelial cells that express a Ca^2+^ biosensor were activated with extracellular ATP, imaged for 13 minutes, and fixed for mRNA abundance quantification using MERFISH. Pairing of cells between the two modalities of the experiment created a unique dataset of 5128 cells with 314 timepoints of Ca^2+^ signaling dynamics and counts for 83 transcripts. This dataset was the basis for the work described here.

New analytical frameworks have emerged for understanding complex dependencies in multimodal measurements with intractable data distributions. Information theory provides a powerful analytical framework for understanding the relationship between system structure and output (Brennan *et al*, 2012). Shannon’s mutual information is a statistical approach for measuring the magnitude of shared, or symmetric, information between two random variables. This framework is powerful because it captures nonlinear relationships and measures true dependence in absolute terms, though it has been difficult to apply to biochemical systems without strict assumptions about the data distribution (Tostevin & ten Wolde, 2010; Uda *et al*, 2013). However, multiomic measurements of single cells often involve different data types that are difficult to relate, that is to define a joint probability distribution. In the case of Ca^2+^ signaling networks, signaling data is sampled from a continuous process whereas RNA abundances are discrete. Many paradigms rely on separate analysis of each data type, often via dimensionality reduction or clustering, before relationships can be quantified (Fang *et al*, 2021; Lee *et al*, 2020; Welch *et al*, 2019). While other approaches (e.g. binning or kernel-density estimation) exist for defining a joint probability distribution over some data types, they fail to perform well outside of strict assumptions about the distributions (e.g. gaussianity) or limited dimensionality; a general, scalable approach is necessary to reduce the need for complex and highly specific analytical pipelines that have emerged (Gayoso *et al*, 2021; Zuo & Chen, 2021). Highly flexible neural networks have demonstrated their ability to estimate characteristics of these probability distributions to allow a deeper understanding of the statistical and information theoretic properties of the data. Deep learning has proven useful for classification of and feature generation from ERK and Akt signaling dynamics (Jacques *et al*, 2021). However, direct interpretation of latent embeddings in these neural network outputs is challenging. An alternative use of deep learning methodologies is a universal functional approximator where neural networks are used to approximate unknown functions to achieve different objective functions. This approach was codified within variational inference and has been proven very useful in probability estimates (Blei et al. 2016). For complex data of mixed types where mutual information is analytically intractable, optimization of neural network functional approximator could be used to find a lower bound. This approach was recently demonstrated under the name Mutual Information Neural Estimator (MINE), which uses a deep neural network to learn a function that can encode the data and find a tight lower bound on the mutual information (Belghazi *et al*, 2018). Briefly, MINE is a universal function approximator that searches for a mapping function *T* in a large space of encoder functions parameterized by *θ* such that *T*_*θ*_: (*G*, *Ca*^2+^) → ℝ. Letting 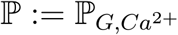 represent the joint probability and 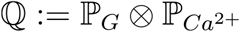 represent the product of the marginals, the mutual information between *G* and *Ca*^2+^ is *I* (*G*, *Ca*^2+^) = *D*_*KL*_(ℙ||ℚ). Using the Donsker-Varadhan representation of the KL divergence, the model parameters *θ* are optimal when gradient ascent has maximized 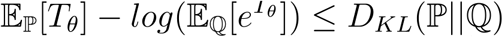 which represents a lower bound on the mutual information. MINE is highly flexible because it makes almost no assumptions about the structure of the data. MINE searches through a large function space for the optimal transformation function to encode the data types assuming there are enough samples to constrain the model. The result is a lower-bound estimate of the mutual information between paired modalities of almost any dimensionality and complexity.

The recent technological development in multiplexed single cell measurements and machine learning approaches for the inference of mutual information could be integrated to provide direct quantification of the phenotypic information content of mRNA abundances. Here, we utilize these developments and focus on a model with timescale separation between an emerging phenotype and mRNA abundance. We relied on highly multiplexed FISH-based quantification of mRNA levels that is more accurate than sequencing based approaches and also allows integration with other imaging modalities. Inference of mutual information was done using the Ca^2+^ signaling network as a model; we fit MINE on various subsets of 83 genes and 314 Ca^2+^ timepoints to quantify the contribution of transcript abundance to signaling dynamics. To establish a baseline, we first calculated the dependency between individual genes and Ca^2+^ signals. We then calculated the mutual information between gene pairs and Ca^2+^ signals to account for redundancy. Gene sets of all sizes were then sampled using various strategies to measure how information changes with set size. Using PCA, we evaluate how useful phenotypic information accounts for transcript-level variance. Overall, we demonstrate a new information theoretic framework for analyzing paired single cell data that provide a quantification of the dependency between sets of mRNAs and an emergent cell-scale dynamic phenotype.

## Results

To investigate the information content of transcript counts and dynamic Ca^2+^ signals, we first analyzed each modality on their own. Ca^2+^ signals display significant heterogeneity across cells (Fig 1A). Likewise, most transcripts had a large range of abundances across cells, though distributions varied depending on the transcript. Pairwise correlation coefficients were calculated for 83 genes across 5128 cells (Fig 1B). The magnitude of the correlations for all gene pairs were relatively low with an average of *r* = 0.16 compared to just cell cycle genes at *r* = 0.44. One interpretation of the low correlations and heterogeneous transcript distributions is that transcripts contain unique information. To test this hypothesis, we calculated the differential entropy of genes using PCA (Fig 1C-E). Because each principal component is an independent, weighted sum of the row vectors of the data, we can approximate the differential entropy among orthogonal components assuming normality via the central limit theorem. Differential entropy across principal components does not measure the information in absolute terms, but can describe how the information is distributed relative to the explained variance. We found that 6 principal components explain 75% of the variance, but only 15% of the entropy. The contrast between Fig 1C and 1D appears contradictory in that few orthogonal components explain most of the variance, yet entropy is steadily added across components with no obvious plateau. This analysis shows that simple measures such as explained variance that are often used for dimensionality reduction are not necessarily appropriate proxies of information content. Accounting for relevant, phenotypic information could help resolve discrepancies between explained variance and differential entropy. To estimate the information content in Ca^2+^ signaling we took advantage of its dynamic patterns. Differential entropy of Ca^2+^ can be estimated using FFT spectral entropy, a scale-invariant measure of information (Burg, 1975). The periodogram shows a continuum of signal frequencies with apparently low variance (Fig 1F). Information can be calculated using this distribution of frequencies, which we found to be 4.2 bits. The mutual information between mRNA abundance and Ca^2+^ signaling is bounded by the distribution with the lowest entropy. Thus, 4.2 bits provides a likely upper bound to the true mutual information between transcripts and Ca^2+^ signals.

**Figure 1.**
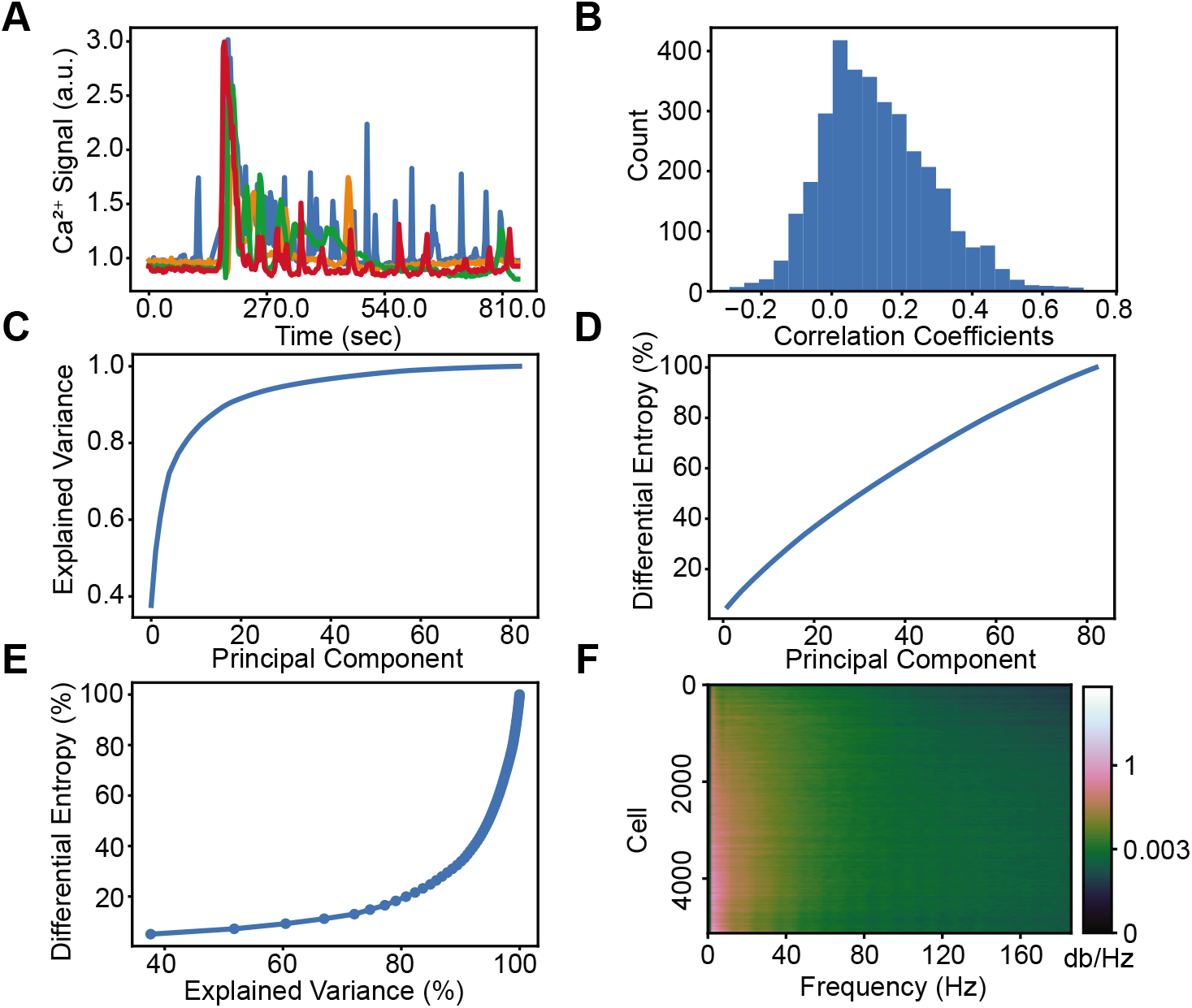
Structure of Gene and Ca^2+^ Data. A) Representative examples of Ca^2+^ dynamics of four cells in the dataset. B) A histogram of the pairwise gene correlation matrix (tri-up) which highlights the relatively low correlations. C) Explained variance of mRNA transcript counts from PCA. D) Differential entropy of transcripts estimated by PCA. E) Plot of explained variance (panel C) vs differential entropy (panel D) with an increasing number of principal components. Collectively, panels C-E show that most of the entropy comes from components that do not explain much of the variance. F) Dynamic Ca^2+^ signal periodogram (cropped to show only the lower wavelength, higher power frequencies). Ca^2+^ dynamic signals were found to contain a spectral entropy of 4.2 bits.

To quantify how useful phenotypic information is distributed across genes, we estimated mutual information between individual genes and Ca^2+^ signals (Fig 2A). Most individual genes contain significant information about Ca^2+^ signals, an average of 0.7 bits, and the most informative genes individually account for 57% of the total mutual information between all genes and Ca^2+^ which is 2.5 +/-0.4 bits. Cell cycle-associated genes were well distributed throughout the list, whereas genes coding for Ca^2+^ and/ or calmodulin-dependent proteins like *PPP3CA* and *CCDC47* were at the top of the list. Interestingly, the sum of the information contained in each gene is significantly larger than the total I(G;Ca^2+^), indicating a high degree of redundancy (Fig 2B). The average mutual information between a single gene and Ca^2+^ signals is 0.7 bits (Fig 2C); the average gene contains about 27% of the signaling information, but how the information is shared across genes is not immediately clear. We further tested whether informative genes, i.e. gene that have high average pairwise mutual information to other genes, are also informative about Ca^2+^ dynamics (Fig 2D). Overall, genes that are more informative about Ca^2+^ signaling are also more informative about the expression of other genes. These genes that are highly informative about Ca^2+^ and many other transcripts may be interpreted as summary genes containing redundant, but distributed information. The second most informative single gene, *PPP1CA*, exemplifies this effect, as it codes for a subunit of PP1 that interacts with >200 regulatory proteins involved in a myriad of critical cell processes. Notably, the top two most informative genes, *PPP3CA* and *PPP1CA*, are both broadly connected phosphatases; kinases and phosphatases were consistently informative and concentrated towards the top of the list. However, from this analysis alone it is not clear how many genes contain redundant information and to what extent.

**Figure 2.**
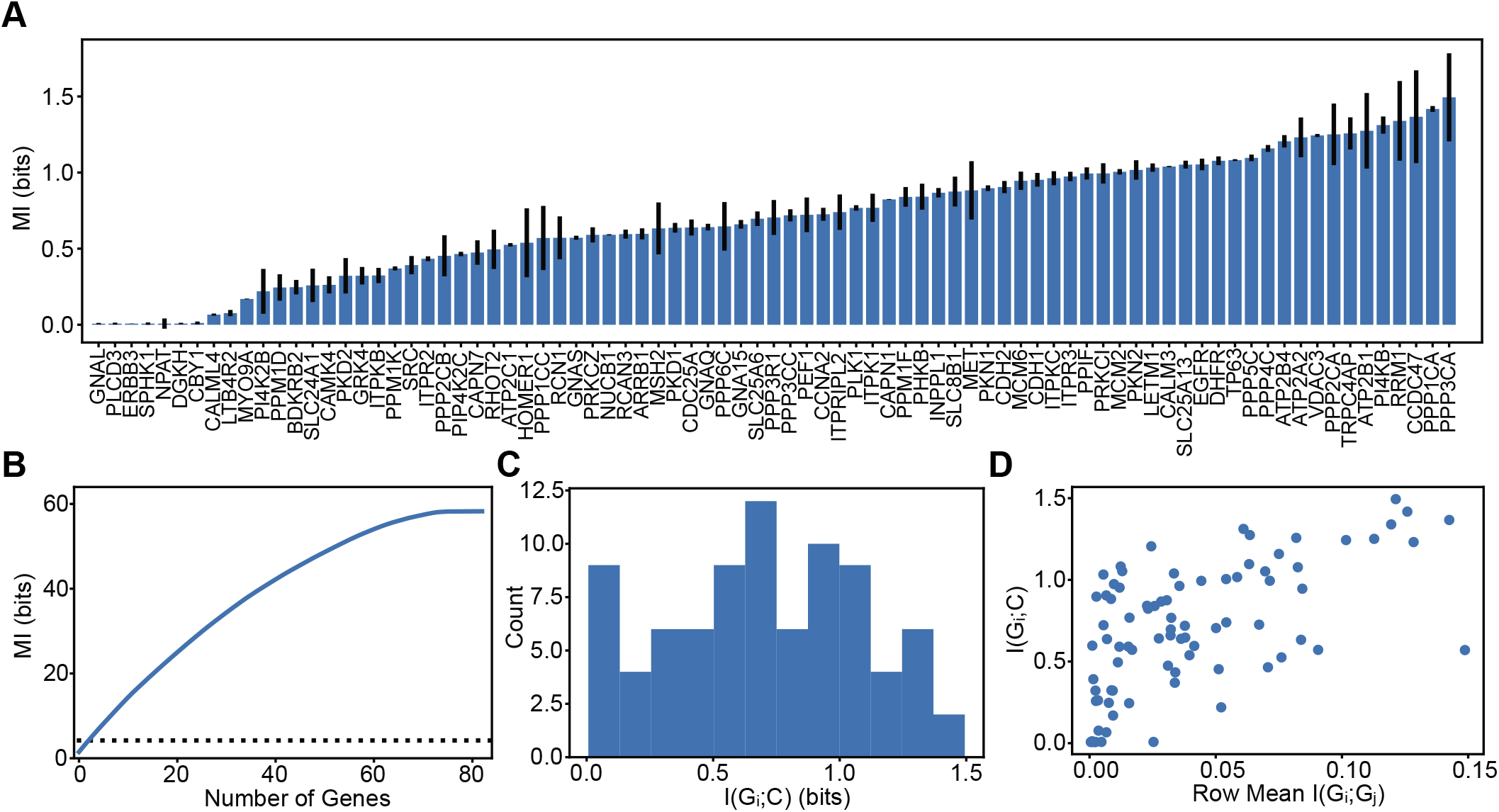
Mutual Information Between Individual Genes and Ca^2+^ Signals. A) I(G_i_;Ca^2+^) sorted from least to greatest. B) The blue line shows the cumulative sum of I(G_i_;Ca^2+^ from (A) sorted from greatest to least, and individual genes appear to contain a lot of information (56 bits) about Ca^2+^ signals. The black dashed line shows mutual information between all 83 genes and Ca^2+^ dynamics estimated to be 2.5 bits. C) Histogram of (A) showing the mean I(G_i_;Ca^2+^) is 0.7 bits. D) I(G_i_;G_j_) represents the pairwise mutual information between genes, the information that genes have about each other. This plot shows that genes that are more informative about other genes tend to be more informative about Ca^2+^ dynamics.

To better understand how the superfluous information in Fig 2B is distributed among genes, we calculated the Synergy Redundancy Index (SRI) between gene pairs with respect to Ca^2+^ (Dietterich *et al*, 2002; Schneidman *et al*, 2003). SRI(G_i_,G_j_ | Ca^2+^) measures the information overlap between genes by subtracting I(G_i_;Ca^2+^) and I(G_j_;Ca^2+^) from I({G_i_,G_j_};Ca^2+^). A gene pair with negative SRI means that the sum of the mutual information between each gene and Ca^2+^ was greater than the gene pair, so the genes must contain some of the same information (redundant). A positive SRI indicates that there is more information about Ca^2+^ in the gene pair than in the sum of the individual genes (synergistic). An SRI of 0 describes a pair of genes that are independent, containing unique and non-overlapping information about Ca^2+^. Calculating SRI between all gene pairs reveals that most pairs are significantly redundant (Fig 3A, B). On average, gene pairs share 0.43 bits which accounts for 61% of the phenotypic information contained in the average individual gene. Further, the more informative a gene is about Ca^2+^, the more redundant it is with other genes (Fig 3C). This finding supports that some genes aggregate information from many others and the more information a gene has, the more it shares. Consistent with findings in Fig 2A, phosphatases and kinases like *PPP3CA*, *PRKCI*, *PPP2CA*, *PI4KB*, and *PPP1CA* were among the most redundant. Interestingly, some genes are highly synergistic on average. One such synergistic gene is *PLCD3*, which appears to have no information about Ca^2+^ on its own, but suddenly becomes informative in gene pairs. *PLCD3* codes for an isoform of phospholipase C, a critical step in the Ca^2+^ in the signal transduction of extracellular ATP. It is surprising that *PLCD3* expression appears to contain little information about Ca^2+^ on its own considering its relevance to stimulus sensing, but this apparent paradox is reconciled by its high synergy. The most synergistic gene on average was *ATP2C1*, which codes for a calcium-transporting ATPase that couples ATP hydrolysis with Ca^2+^ transport into the Golgi lumen. Genes which are critical for modulating Ca^2+^ concentrations in the cytoplasm represent steps in linear processes, rather than cooperating with many other components to achieve their function. Generally, the most synergistic genes were not very informative on their own (Fig 2A), but became informative in a group of 2 (Fig 3C). The high degree of synergy suggests that these genes provide contextual or conditional information that is absent from most other genes, even genes that were independently informative. Although, genes typically function in larger sets beyond pairs, and a thorough understanding of transcriptional information requires evaluation of higher order interactions.

**Figure 3.**
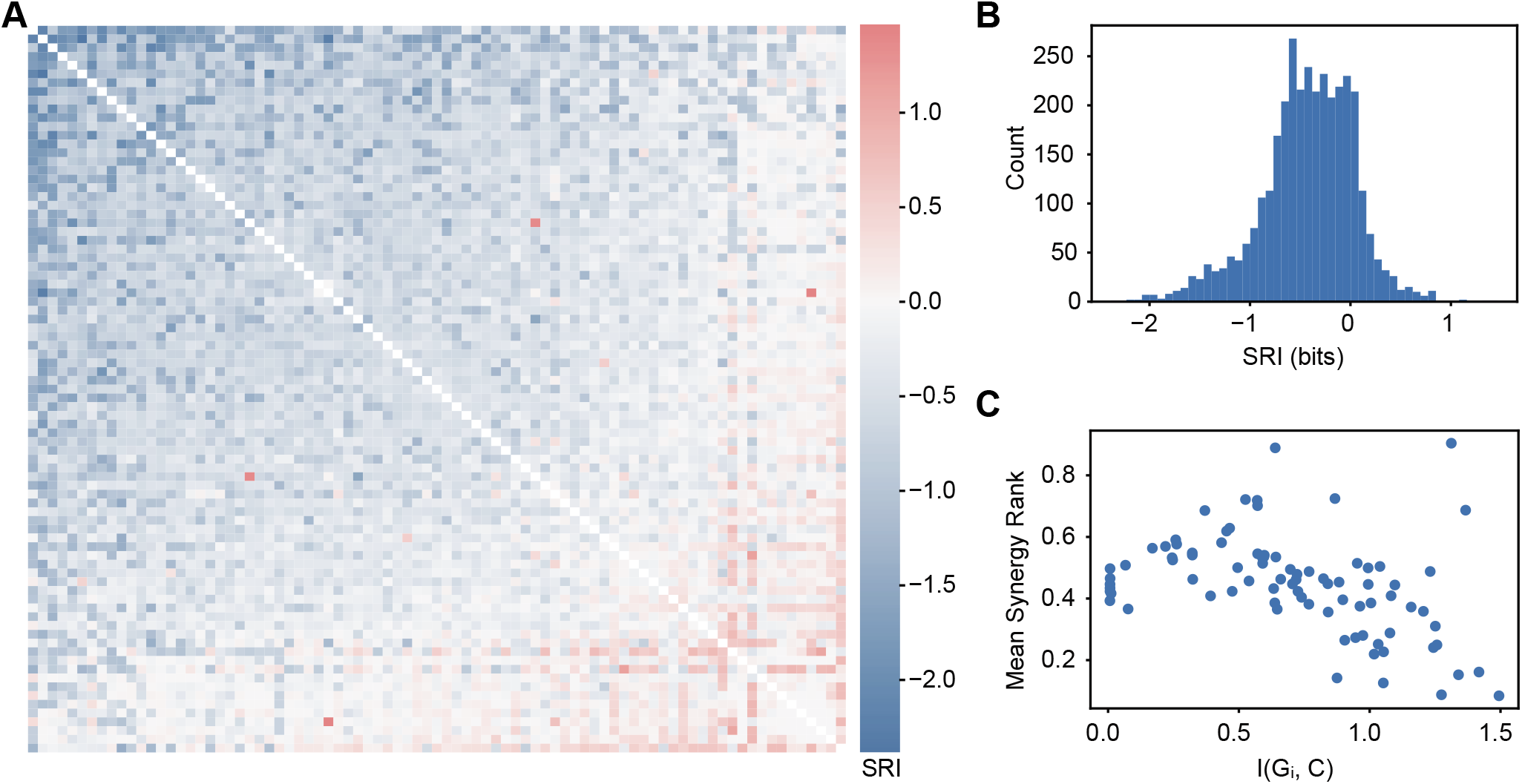
Synergy and Redundancy of Gene Pairs With Respect to Ca^2+^. A) SRI(G_i_,G_j_|C) sorted by average SRI. B) Histogram of SRI showing that most gene pairs are highly redundant with an average score of −0.43 bits. C) The mean rank of all synergistic pairs compared to the mutual information between that gene and Ca^2+^ signals, (spearman *r*=0.5, *P*<2e-6), indicating that genes with more information about Ca^2+^ are also more redundant.

To explore the mutual information between Ca^2+^ and gene sets of various sizes, we tested various sets using gene annotations, a sequential search, and PCA. To quantify set level information at a functional level, we summarized pairwise SRIs based on gene annotations (Fig 4A). Calculating the mean SRI for combinations of annotations revealed how different functional gene sets contain phenotypic information. Pairs within an annotation are more redundant than between annotations with an average difference of ~0.1 bits.The Ca^2+^/ER annotation contains the most redundancies by a large margin, whereas the miscellaneous category “Other” is the most synergistic which can be explained by the functional diversity in this group. The Ca^2+^/ER annotation contains the genes most relevant to the stimulus and appear to provide similar information. To understand how phenotypic information depends on gene set size, we calculated the mutual information between Ca^2+^ signals and gene sets of all sizes. Because testing all possible sets is prohibitively computationally expensive, we first sampled random sets of all possible sizes (4B). Each set size was sampled 4 times. For random sets, 53 genes contained 90% of the phenotypic information. To understand the upper and lower bounds on information in each set, we performed two directed heuristic searches. The directed searches first picked the most (least) informative gene, and then tested every possible addition to the set to add the member that contributed the most (least) information until the sets were of maximum size. The upper bound in green shows that the information quickly plateaus as the best 12 genes contain 90% of the phenotypic information, and all further additions contribute minimal additional information. The lower bound in purple shows the unique information per gene given the set, sorted from least to greatest. Because the lower bound always adds the least informative and most redundant genes first, the last genes contain the most unique information. *PPP3CA* is the first gene added to the upper bound and the last gene to be added to the lower bound, which means it must have both the most absolute information and the most unique information. Interestingly, the growth of information in the least informative set was approximately linear, meaning that there is always some unique information in every gene. The slope of the lower bound is 0.03 bits, which represents the average unique information per gene.

**Figure 4:**
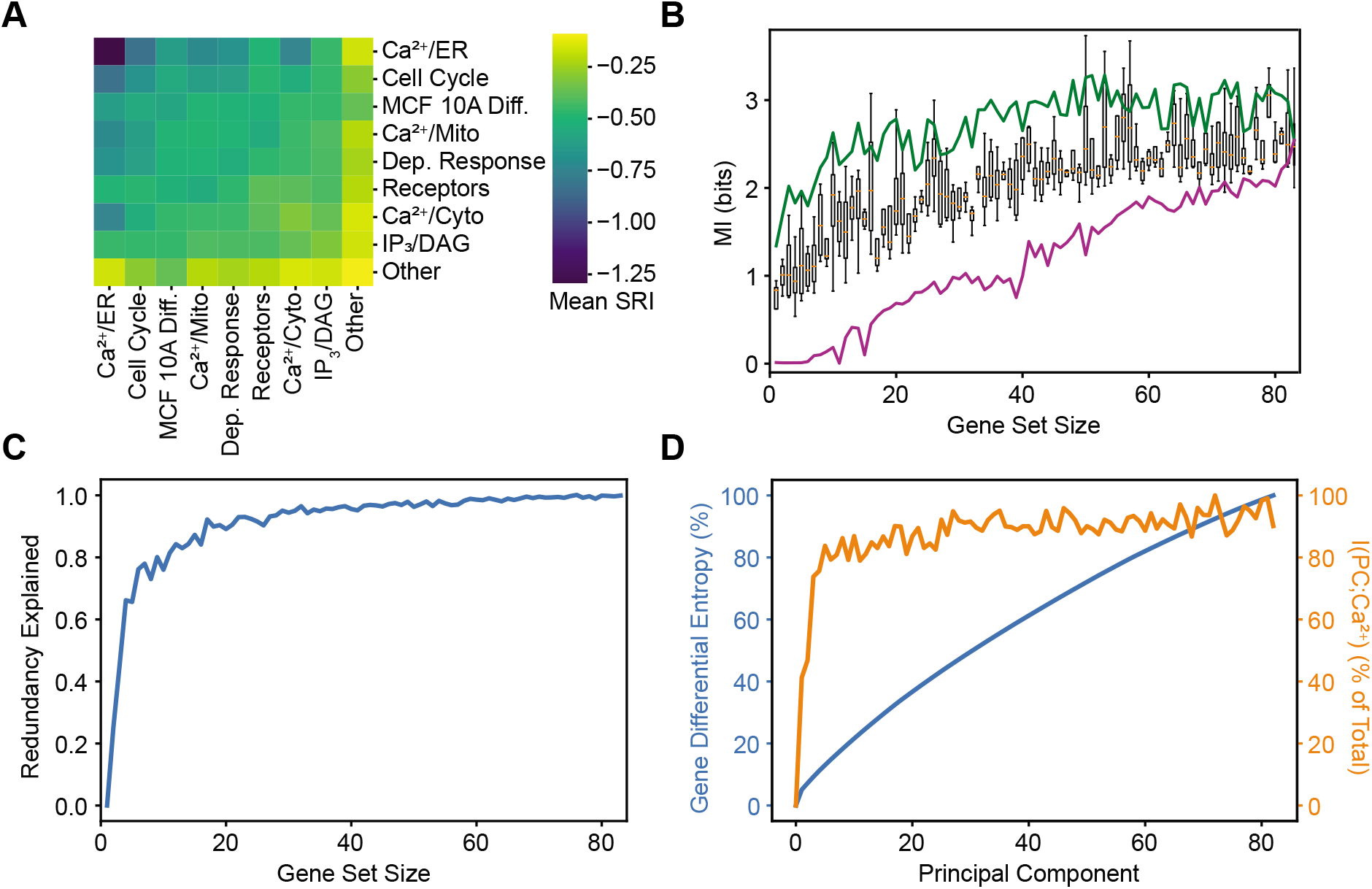
Mutual Information Between Gene Sets and Ca^2+^ Signals. A) Mean pairwise SRI from Figure 3 for sets based on annotation. MCF 10A differentiation and Ca^2+^ dependent response are abbreviated. B) Gene sets of various sizes were constructed using three different strategies: an upper bound (green) that always adds the most informative gene to the set given the genes already included, random strategy (boxes) that samples random sets of genes, and a lower bound (purple) that always adds the least informative gene to the set given the genes already included. C) The blue line shows the fraction of redundant information using the expected value of I({G_0_, …, G_n_};Ca^2+^) from Equation 3. D) A y-y plot of gene differential entropy in blue (same as Figure 1D) and the mutual information between gene principal components and Ca^2+^ in orange. Both values are normalized by their respective max values.

Using the mean mutual information between a gene set and Ca^2+^, we can also estimate the “redundancy explained” of all sets of genes of a given size according to Equation 3 (Fig 4C). We found that sets of only three genes explain 66% of the redundancy. Small gene sets contain much more redundant phenotypic information than larger gene sets. The point at which this curve begins to level off can be interpreted as a fundamental set size above which most phenotypic information lies within the sets. We observe that small gene sets contain most of the information on Ca^2+^ dynamics suggesting that higher order interactions in larger sets are not required to capture the full dependency between mRNA abundance and Ca^2+^ dynamics.

Finally, we calculated the mutual information between transcript principal components and Ca^2+^ signals to compare with differential entropy and understand how useful phenotypic information is distributed (Fig 4D). In agreement with Fig 4C, phenotypic information saturates quickly with only 3 principal components accounting for 74% of the total mutual information between transcripts and signals. This result starkly contrasts with the differential entropy of gene principal components independent of Ca^2+^ signals which rises slowly and does not appear to plateau. By accounting for phenotypic information, far fewer orthogonal components are required to preserve the useful information. The difference between these curves indicates that focusing on phenotypic information may filter or compress transcriptional information. I(PC;Ca^2+^) resembles the curve in Fig 1C, though still plateaus more quickly. These results confirm that phenotypic information is mostly explainable by a few components and higher order interactions do not significantly contribute.

## Discussion

The complexity of biological regulation is staggering. While many details about biological networks are known, the gaps in our knowledge make some simple questions very challenging to answer. For example, to what degree does abundance of one set of molecules matter? Specifically, does the abundance of mRNAs matter for the regulation of complex cellular phenotypes such as signaling response to a ligand? Here we provide a framework for answering such questions through the combination of paired single cell data and application of recent advances at the interface between machine learning and information theory (Belghazi *et al*, 2018). Applying a recently developed framework for mutual information estimation to single cell data of multiplexed mRNA levels paired with live cell imaging allowed us to quantify the strength of the causal connection between mRNAs and Ca^2+^ signaling. We found that approximately 60% of Ca^2+^ signal information exists in transcript counts, which is 2.5 (+/− 0.4) bits. Furthermore, the framework we developed provides key information about information synergy and redundancy, can be used to quantify the joint information in sets of genes, and reveals how overall dependency changes with the size of the set. On average, genes were found to contain 61% redundant information with each other, though nearly all genes contained some unique information. Genes that appeared to contain little phenotypic information individually were in fact the most synergistic and became informative in pairs. The unique information present in gene sets is best visualized in Fig. 4B, which illustrates the difference in information among the most, least, and average set. In the best case, only 12 genes contain 90% of signal information, which is significantly fewer than an equally informative 53 random genes. While all genes contain unique information, some sets are still significantly more informative than others likely due to their role in the signaling network. Decomposition by principal components (Fig. 4D) revealed a rapid plateau in phenotypic information, starkly contrasting the increasing growth in differential entropy. These results demonstrate the utility of information theoretic analysis in quanityfing the phenotypic information of mRNA abundance.

The framework we propose is very general and can be applied to any two “slices” within a complex biological regulatory network. Our numerical experiments (Supplementary Material) demonstrate that with minor adaptations for bias removal, MINE can robustly estimate mutual information between two high dimensional vectors containing 100+ features. The generalizability of this framework provides a new tool to put weights and interpretable numbers on different “arrows” within complex biological regulatory networks. Importantly, such “arrows” do not necessarily represent direct mechanistic steps. There are numerous reactions that occur post transcriptionally to determine Ca^2+^ signaling responses. Yet, using MINE we were able to infer the individual contribution of each gene in controlling the emergent phenotypes. Furthermore, through the use of pairs of genes and estimation of the effect of gene set size, we determined how information between multiple mRNA types is integrated. This inference showed that despite the information having to propagate through multiple layers of regulation, it still shows significant dependency. Even though correlations between mRNA and protein levels are generally low, phenotypically relevant information is still preserved in the transcriptome. Our results supports the use of mRNA measurements to infer useful phenotypic characteristics of cell populations. An important feature of our analysis is that all inference was done relaying on natural heterogeneity without any experimental perturbation to gene expression circumventing compensation and non-linear dependencies that are common pitfalls of perturbation analysis (Welf & Danuser 2014).

While the framework we propose is very general, our findings are systems specific and will change depending on the set of genes and measured phenotypes. Here we focused on Ca^2+^ signaling in response to activation of GPCR in a clonal population of MCF 10A cells. In previous work, we estimated that a cellular population is composed of multiple subtypes (Yao *et al*, 2016) and have shown that mRNA variability is dominated by cell state differences with a minor contribution from transcriptional bursting (Foreman & Wollman, 2020). Our current finding that 60% of information in the emerging Ca^2+^ signaling phenotypes can be attributed to cellular transcriptional state largely agrees with these previous findings. It is likely that in other systems, decomposition of information content will differ from the 60% transcriptional and 40% post transcriptional measured here. For example, broad phenotypes such as cell type classification that often correspond to larger and highly patterned transcriptional differences will likely show higher levels of transcriptional dependency.

Our approach has several limitations, experimental and computational, that will need to be addressed in future work. Experimentally, gene selection, i.e. the expression of which genes are measured, is limited due to gene length, specific sequence, and other experimental constraints that are continuously improving. Furthermore, the approach could be applied to tissue samples with much higher population diversity where the relationship between transcripts and phenotype is more relevant. Computationally, because of stochastic gradient ascent, the model’s estimates are somewhat noisy and required multiple replicates. Additionally, we were limited to exploring the effect of set sizes with search strategies and only exhaustively examined pairwise dependencies because the model was computationally expensive to run. None of the search strategies are guaranteed to find the truly most or least informative set because doing so would require a prohibitively time consuming exhaustive search.

Despite these limitations, MINE was able to provide an interpretable and scalable quantification of dependency between transcript sets and Ca^2+^ signaling.

Recent advances in single cell technologies are making high-dimensional, multimodal measurements feasible. Statistical descriptions of complex phenotypes will become increasingly useful as single cell experiments generate more multimodal and multiomic data. Integrating multiple different data types is still a challenge in the field, and this work represents a new approach to synthesize statistical descriptions of high-dimensional, multimodal data that does not make any assumptions about the underlying functional relationships. This unbiased approach will enable a deeper understanding of complex, multidimensional data by quantifying the dependency between any single cell phenomena.

## Methods

### Data Selection

Data collection is described in previous work (Foreman & Wollman, 2020). Of the 336 genes measured, 150 genes were measurably expressed and the top 83 were chosen by the highest magnitude z scores from multiple linear regression.

### Preprocessing

Transcript counts from the 83 genes and 314 timepoints of Ca^2+^ signals were independently z-score normalized. Normalization was applied to the entire matrix of all cells for each data type (e.g. 5128×83 for transcripts) and not to individual columns, preserving relative magnitude across genes and timepoints.

### MINE

Hyperparameters were chosen by fitting analytically tractable data from an additive white gaussian noise model of the data across a range of strengths of dependence (SF1). Additional bias correction was implemented by fitting *I*_*obs*_(*t*) = *I*_*true*_(1 - *a*) ∗ *e*^−*b*∗*t*^ + *ct*, where *I*_*true*_, *a*, *b*, and *c* are the fitting parameters and is the number of iterations (SF2). Convergence tests were performed on the real data by comparing the residuals of the bias correction fit. The chosen hyperparameters of 600 hidden units and a learning rate = 3e-4 resulted in the highest yield, i.e. fewest failed fits.

### Synergy Redundancy Index (SRI)

The Synergy Redundancy Index was developed to evaluate information about a stimulus shared among a small population of cells (Dietterich *et al*, 2002). Equation 1 describes the calculation, which involves comparing pairwise and individual mutual information between genes and Ca^2+^ signals.

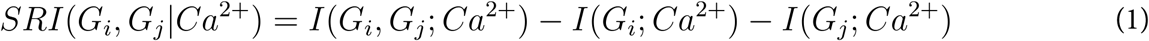

First, the mutual information between each unique pair of genes and Ca^2+^ were estimated, I(G_i_,G_j_; Ca^2+^). Then I(G_i_; Ca^2+^) was calculated, and Equation 1 was calculated for all genes.

### Redundancy Explained

This metric represents the amount of extra information assuming no redundancy between elements. Equation 2 uses an expected value multiplied by the number of sets to calculate the non-redundant information (NRI) as if all individual sets contain unique information:

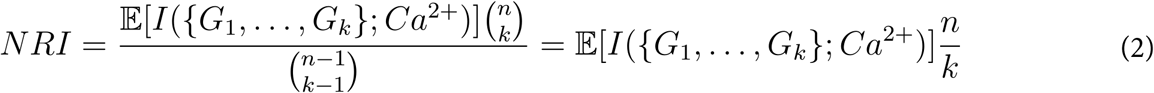

k = set size, n = total number of genes

From Equation 2, we can calculate the redundancy explained (Equation 3):

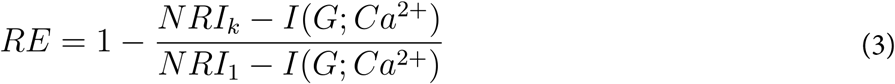

## Acknowledgements

We thank Robert Foreman for significant analytical and conceptual contributions, as well as Alon Oyler-Yaniv for suggesting spectral entropy to quantify Ca^2+^ signal information. We gratefully acknowledge the support of NVIDIA Corporation for donation of the two Titan V GPUs used for this research.

## Supplementary Materials

MINE estimates mutual information by calculating the KL Divergence between the marginal distributions and the joint distribution. This KL divergence represents the distance between these distributions, which is nonnegative such that 0 represents complete independence. MINE uses the Donsker-Varadahn representation of the KL Divergence to evaluate a function that maps samples of the data to the set of all real numbers. Gradient ascent finds the parameterization of this function which maximizes the mutual information for a tight lower bound.

To validate MINE’s estimates before applying it to the real data, we evaluated MINE on multivariate gaussian distributions with an analytically calculable mutual information between toy genes and toy Ca^2+^ (I(TG;TC)). The “toy” data was created using a standard additive white gaussian noise (AWGN) model with tunable dependency, entropy, and dimensionality (SF 1). A multivariate gaussian distribution (5128,16) was defined based on a specified covariance structure (SF 1B). The covariance of the distribution is somewhat arbitrary; it is only required that the matrix is invertible and tunable over a range of dependencies. The covariance structure is shown as 4 symmetrical quadrants in the matrix: the off diagonal quadrants are −α and the on-diagonal quadrants are α (SF 1A). Toy signals were created by applying a tunable amount of noise to the toy genes (SF 1C).

**Supplementary Figure 1.**
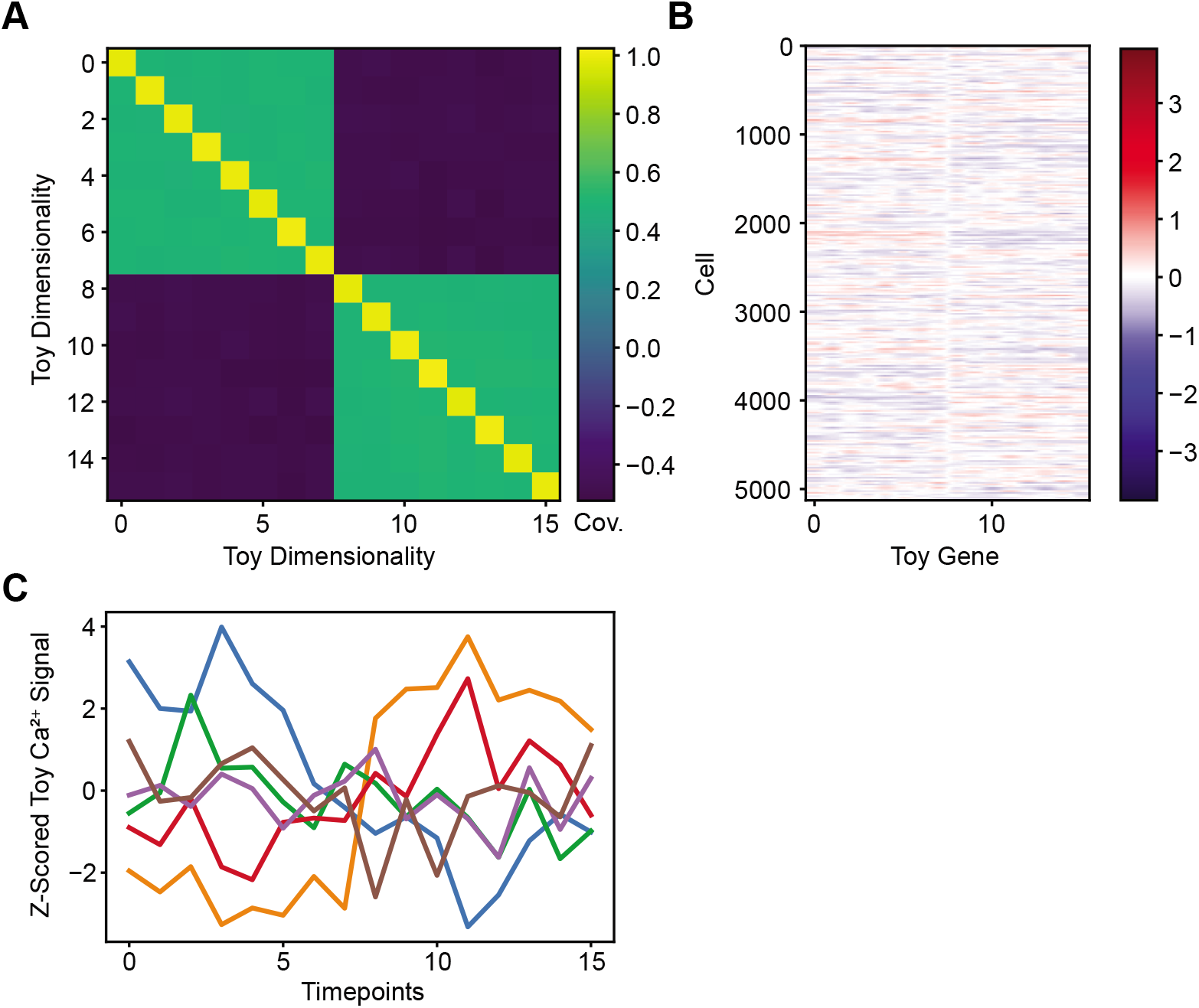
Additive White Gaussian Noise (AWGN) Toy Model. A) Covariance matrix for toy data. Quadrant values are determined by a hyperparameter α, where off-diagonal quadrants are set to −α and on-diagonal quadrants are set to α, with variance set to 1. B) Toy genes are determined by the covariance matrix. C) Toy signals are calculated by adding a tunable amount of noise to the toy gene matrix: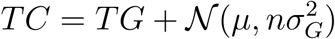.

Evaluating MINE on the toy data, we found that the model occasionally produces a non-converging, biased mutual information estimate that increases linearly with training (SF 2A). To quantify and remove this bias, we fit MINE’s output to a statistical model: *I*_*obs*_(*t*) = *I*_*true*_(1 - *a*) ∗ *e*^−*b*∗*t*^ + *ct*, where *I*_*true*_, *a*, *b*, and *c* are the fitting parameters and *t* is the number of iterations. Finding the optimal *I*_*true*_ factors out the linear bias *c* and produces a converging and accurate estimate of the analytically determined I(TG;TC) (SF 2B). The mean over the three replicates yields a final slope of 5.8e-6, showing reliable, asymptotic convergence to the true mutual information value. Applying this solution allowed rigorous definition of a convergence criterion that adaptively determines when training can conclude (SF 2C). This criterion uses moving averages to determine convergence, and expectedly results in higher residuals for fits with fewer iterations (SF 2D). Stricter convergence thresholds can fail to converge because of noise in the output; 1.4 was chosen as the final threshold due to its high yield, fast convergence, and low mean residual (SF 2E). In addition to the low residuals, we also found a pearson correlation of 0.97 across a range of I(TG;TC) (SF 2F). These results support the use of our bias-corrected MINE with the chosen hyperparameters.

**Supplementary Figure 2.**
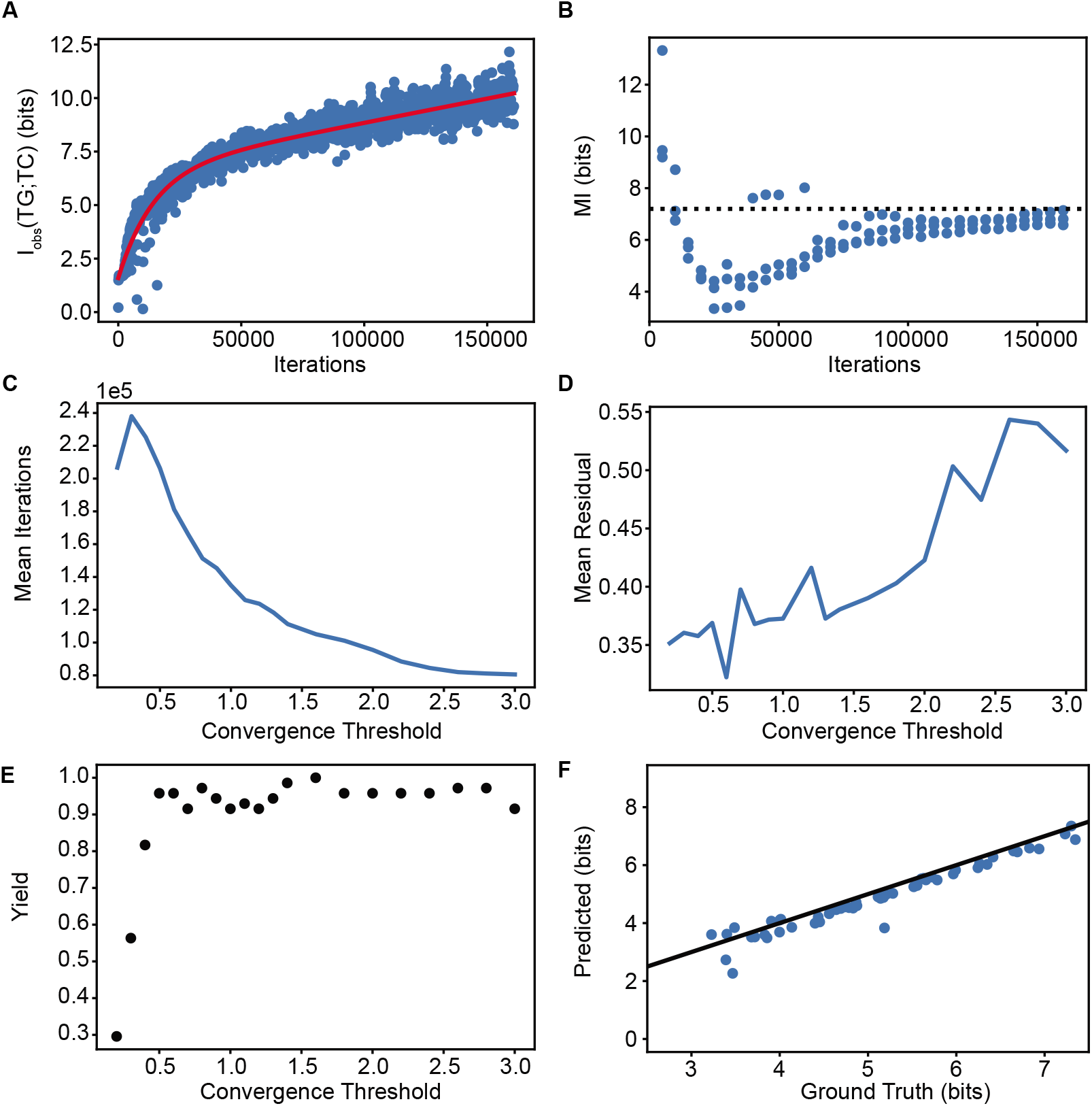
Fitting Model on Toy Data. A) Blue points are the raw output from MINE. The model occasionally failed to converge at the analytical I(TG;TC) value, as shown. Bias correction fit using BFGS optimization is shown in red. B) Blue points show the bias correction estimate of the true I(TG;TC) during training from 3 replicates. The dotted black line shows the true, analytical I(TG;TC) value. C) An adaptive convergence criterion based on the difference between moving averages was used to decrease training time. Higher convergence thresholds, i.e. larger differences between moving averages, result in fewer iterations. D) Faster convergence expectedly results in poorer fits with higher residuals on average. E) If the convergence threshold is too stringent, the optimization algorithm may be unable to find a suitable fit for bias correction. The yield was calculated over many toy datasets with a range of parameter values. F) Toy models with a large range of analytical mutual information values were fit using bias-corrected MINE, pearson *r* = 0.97.

We then applied bias-corrected MINE to the full data. We found that the bias correction works well on the real data and eventually converges on a single value (SF 3A). To verify that the results are not limited by sample size, we performed a jackknife correction (SF 3B). The jackknife extrapolation to infinite sample size yielded a result well within the error bars of the estimate at the full sample size. This result indicates that sample size is not limiting.

**Supplementary Figure 3.**
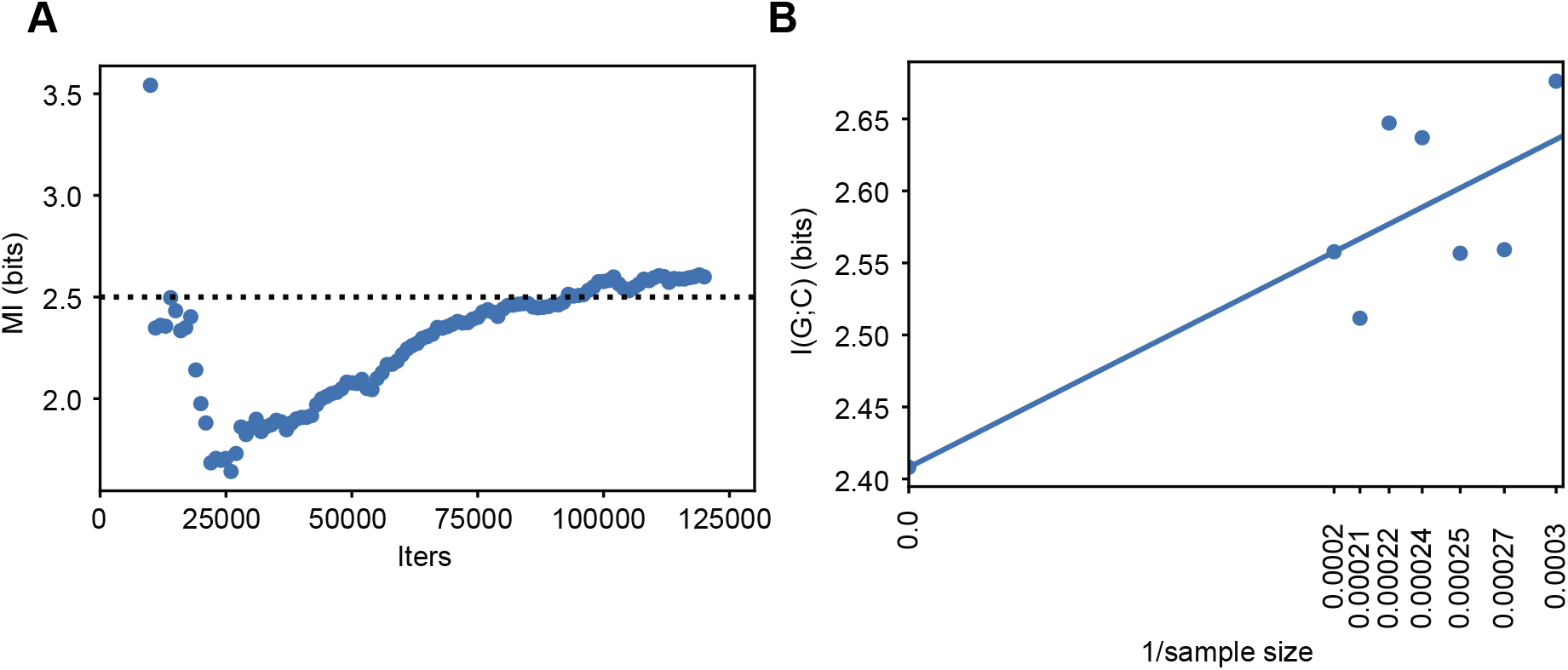
Bias-Corrected MINE on Real Data. A) Full data was fit using bias-correction and the estimate of I(G;Ca^2+^) is shown in blue. The dotted black line is the mean of several samples. B) Jackknife of the data with 7 sample sizes ranging from 3369 to 5128 (all cells). The intercept shown is the extrapolation at infinite sample size, which is well within the estimate of 2.5 ± 0.4 bits.

To estimate the upper bound on the MI, we used FFT spectral entropy. This calculation begins by creating an FFT periodogram (Fig 1D). The Shannon entropy of the resulting distribution of power spectral densities represents the spectral entropy. To verify that spectral entropy produces an invariant measurement of entropy unlike other differential entropy metrics, we first applied various transformations to the signals. These transformations did not change the final entropy, whereas smoothing by convolution did degrade the entropy, as expected (SF 4A). Furthermore, we applied linear interpolations to change the dimensionality and found that most of the entropy comes from low frequencies which are well preserved even with dramatic reduction in timepoints (SF 4B). Increasing the dimensionality by interpolation did not change the entropy estimate. Therefore, we conclude that spectral entropy is a robust, invariant measure of signal information.

**Supplementary Figure 4:**
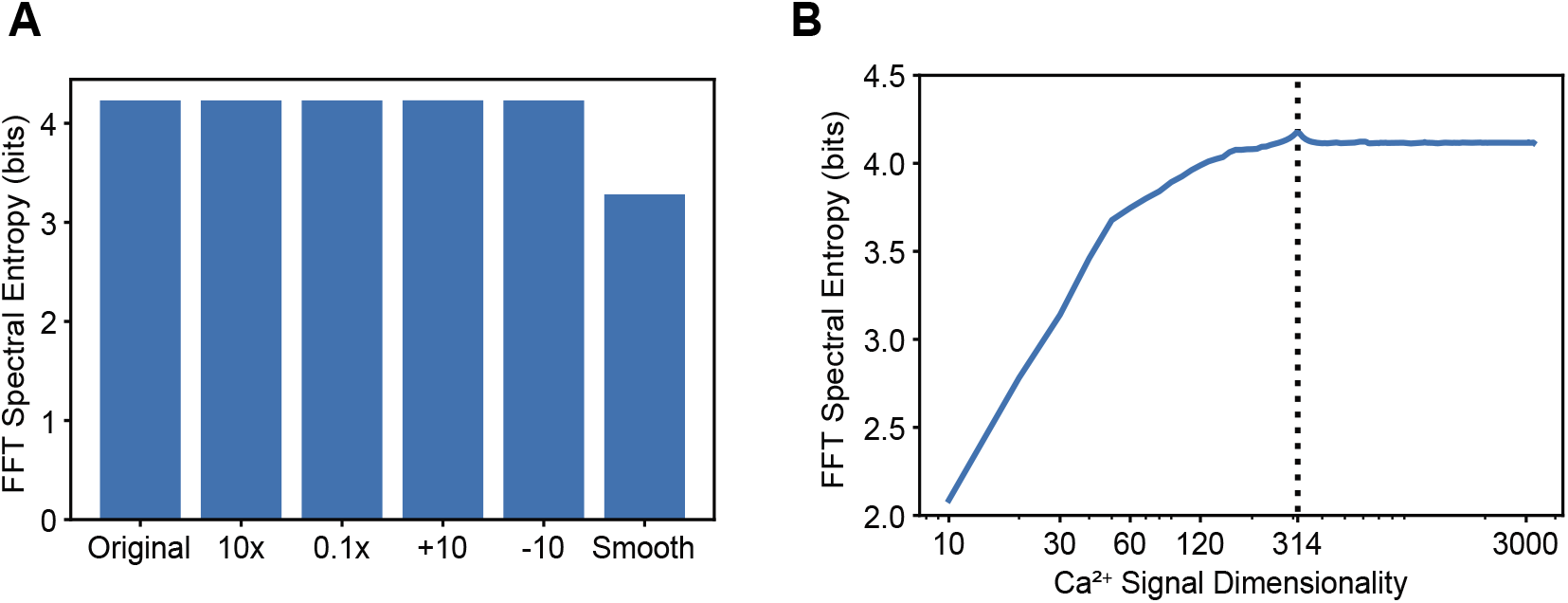
FFT Spectral Entropy Robustness. A) FFT spectral entropy is robust to translations and rescaling; unlike other measures of differential entropy, it does not depend on the scale of the data. Smoothing the data by convolution expectedly results in a loss of information due to removal of high frequencies. B) The dotted black line shows the true dimensionality and the blue line shows spectral entropy as a function of dimensionality. FFT spectral entropy is robust to interpolations that change the dimensionality of the data that do not affect the distribution of frequencies. Expectedly, reducing dimensionality results in a loss of high frequency information, though most of the information is at relatively low or mid frequencies.

To estimate the amount of extra information assuming no redundancy between elements, we calculated the NRI as a function of gene set size. For I(G_i_;Ca^2+^), the NRI is simply defined as 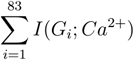 because each gene occurs only once. This equation can be generalized to gene sets of any size by dividing I({G_0_, …, G_n_};Ca^2+^) by the number of times a gene appears in a particular set. Replacing the sum with an expected value multiplied by the number of sets is also useful to avoid having to calculate the value of each element. Equation 2 defines the generalization and is used to calculate the redundancy explained in Equation 3, which represents the fraction of redundant information at a given gene set size.

## References

Adelaja A, Taylor B, Sheu KM, Liu Y, Luecke S & Hoffmann A (2021) Six distinct NFκB signaling codons convey discrete information to distinguish stimuli and enable appropriate macrophage responses. Immunity 54: 916–930.e7

Alves M, Beamer E & Engel T (2018) The Metabotropic Purinergic P2Y Receptor Family as Novel Drug Target in Epilepsy. Front Pharmacol 9: 193

Azeloglu EU & Iyengar R (2015) Signaling networks: information flow, computation, and decision making. Cold Spring Harb Perspect Biol 7: a005934

Bagur R & Hajnóczky G (2017) Intracellular Ca2+ Sensing: Its Role in Calcium Homeostasis and Signaling. Mol Cell 66: 780–788

Balázsi G, van Oudenaarden A & Collins JJ (2011) Cellular decision making and biological noise: from microbes to mammals. Cell 144: 910–925

Belghazi MI, Baratin A, Rajeswar S, Ozair S, Bengio Y, Courville A & Devon Hjelm R (2018) MINE: Mutual Information Neural Estimation. arXiv [csLG]

Brennan MD, Cheong R & Levchenko A (2012) Systems biology. How information theory handles cell signaling and uncertainty. Science 338: 334–335

Burg JP (1975) Stanford University.

Cheong R, Rhee A, Wang CJ, Nemenman I & Levchenko A (2011) Information transduction capacity of noisy biochemical signaling networks. Science 334: 354–358

Darmanis S, Gallant CJ, Marinescu VD, Niklasson M, Segerman A, Flamourakis G, Fredriksson S, Assarsson E, Lundberg M, Nelander S, et al (2016) Simultaneous Multiplexed Measurement of RNA and Proteins in Single Cells. Cell Rep 14: 380–389

Dietterich TG, Becker S & Ghahramani Z (2002) Advances in Neural Information Processing Systems: Proceedings of the 2001 Conference MIT Press

El-Samad H (2021) Biological feedback control-Respect the loops. Cell Syst 12: 477–487

Emert BL, Cote CJ, Torre EA, Dardani IP, Jiang CL, Jain N, Shaffer SM & Raj A (2021) Variability within rare cell states enables multiple paths toward drug resistance. Nat Biotechnol 39: 865–876

Fang R, Preissl S, Li Y, Hou X, Lucero J, Wang X, Motamedi A, Shiau AK, Zhou X, Xie F, et al (2021) Comprehensive analysis of single cell ATAC-seq data with SnapATAC. Nat Commun 12: 1337

Foreman R & Wollman R (2020) Mammalian gene expression variability is explained by underlying cell state. Mol Syst Biol 16: e9146

Gayoso A, Steier Z, Lopez R, Regier J, Nazor KL, Streets A & Yosef N (2021) Joint probabilistic modeling of single-cell multi-omic data with totalVI. Nat Methods 18: 272–282

Genshaft AS, Ziegler CGK, Tzouanas CN, Mead BE, Jaeger AM, Navia AW, King RP, Mana MD, Huang S, Mitsialis V, et al (2021) Live cell tagging tracking and isolation for spatial transcriptomics using photoactivatable cell dyes. Nat Commun 12: 1–15

Gong H, Wang X, Liu B, Boutet S, Holcomb I, Dakshinamoorthy G, Ooi A, Sanada C, Sun G & Ramakrishnan R (2017) Single-cell protein-mRNA correlation analysis enabled by multiplexed dual-analyte co-detection. Sci Rep 7: 2776

Hafner A, Stewart-Ornstein J, Purvis JE, Forrester WC, Bulyk ML & Lahav G (2017) p53 pulses lead to distinct patterns of gene expression albeit similar DNA-binding dynamics. Nat Struct Mol Biol 24: 840–847

Jacques M-A, Dobrzyński M, Gagliardi PA, Sznitman R & Pertz O (2021) CODEX, a neural network approach to explore signaling dynamics landscapes. Mol Syst Biol 17: e10026

Kinnunen PC, Luker KE, Luker GD & Linderman JJ (2021) Computational methods for characterizing and learning from heterogeneous cell signaling data. Current Opinion in Systems Biology 26: 98–108

Lane K, Van Valen D, DeFelice MM, Macklin DN, Kudo T, Jaimovich A, Carr A, Meyer T, Pe’er D, Boutet SC, et al (2017) Measuring Signaling and RNA-Seq in the Same Cell Links Gene Expression to Dynamic Patterns of NF-κB Activation. Cell Syst 4: 458–469.e5

Lee J, Hyeon DY & Hwang D (2020) Single-cell multiomics: technologies and data analysis methods. Exp Mol Med 52: 1428–1442

Liu M, Yang B, Hu M, Radda JSD, Chen Y, Jin S, Cheng Y & Wang S (2021) Chromatin tracing and multiplexed imaging of nucleome architectures (MINA) and RNAs in single mammalian cells and tissue. Nat Protoc 16: 2667–2697

Liu Y, Beyer A & Aebersold R (2016) On the Dependency of Cellular Protein Levels on mRNA Abundance. Cell 165: 535–550

Macaulay IC, Ponting CP & Voet T (2017) Single-Cell Multiomics: Multiple Measurements from Single Cells. Trends Genet 33: 155–168

Mair F, Erickson JR, Voillet V, Simoni Y, Bi T, Tyznik AJ, Martin J, Gottardo R, Newell EW & Prlic M (2020) A Targeted Multi-omic Analysis Approach Measures Protein Expression and Low-Abundance Transcripts on the Single-Cell Level. Cell Rep 31: 107499

Moffitt JR, Hao J, Wang G, Chen KH, Babcock HP & Zhuang X (2016) High-throughput single-cell gene-expression profiling with multiplexed error-robust fluorescence in situ hybridization. Proc Natl Acad Sci U S A 113: 11046–11051

Myers PJ, Lee SH & Lazzara MJ (2021) Mechanistic and data-driven models of cell signaling: Tools for fundamental discovery and rational design of therapy. Current Opinion in Systems Biology 28: 100349

Nelson DE, Ihekwaba AEC, Elliott M, Johnson JR, Gibney CA, Foreman BE, Nelson G, See V, Horton CA, Spiller DG, et al (2004) Oscillations in NF-kappaB signaling control the dynamics of gene expression. Science 306: 704–708

Park S & Pardalos PM (2021) Deep Data Density Estimation through Donsker-Varadhan Representation. arXiv [csLG]

Perkins TJ & Swain PS (2009) Strategies for cellular decision-making. Mol Syst Biol 5: 326

Purvis JE & Lahav G (2013) Encoding and decoding cellular information through signaling dynamics. Cell 152: 945–956

Putney JW (2012) Calcium signaling: deciphering the calcium-NFAT pathway. Curr Biol 22: R87–9

Schneidman E, Bialek W & Berry MJ 2nd (2003) Synergy, redundancy, and independence in population codes. J Neurosci 23: 11539–11553

Schulz D, Zanotelli VRT, Fischer JR, Schapiro D, Engler S, Lun X-K, Jackson HW & Bodenmiller B (2018) Simultaneous Multiplexed Imaging of mRNA and Proteins with Subcellular Resolution in Breast Cancer Tissue Samples by Mass Cytometry. Cell Syst 6: 25–36.e5

Selimkhanov J, Taylor B, Yao J, Pilko A, Albeck J, Hoffmann A, Tsimring L & Wollman R (2014) Systems biology. Accurate information transmission through dynamic biochemical signaling networks. Science 346: 1370–1373

Shaffer SM, Emert BL, Reyes Hueros RA, Cote C, Harmange G, Schaff DL, Sizemore AE, Gupte R, Torre E, Singh A, et al (2020) Memory Sequencing Reveals Heritable Single-Cell Gene Expression Programs Associated with Distinct Cellular Behaviors. Cell 182: 947–959.e17

Spencer SL, Gaudet S, Albeck JG, Burke JM & Sorger PK (2009) Non-genetic origins of cell-to-cell variability in TRAIL-induced apoptosis. Nature 459: 428–432

Stoeckius M, Hafemeister C, Stephenson W, Houck-Loomis B, Chattopadhyay PK, Swerdlow H, Satija R & Smibert P (2017) Simultaneous epitope and transcriptome measurement in single cells. Nat Methods 14: 865–868

Subramanian I, Verma S, Kumar S, Jere A & Anamika K (2020) Multi-omics Data Integration, Interpretation, and Its Application. Bioinform Biol Insights 14: 1177932219899051

Taniguchi Y, Choi PJ, Li G-W, Chen H, Babu M, Hearn J, Emili A & Xie XS (2010) Quantifying E. coli proteome and transcriptome with single-molecule sensitivity in single cells. Science 329: 533–538

Tostevin F & ten Wolde PR (2010) Mutual information in time-varying biochemical systems. Phys Rev E Stat Nonlin Soft Matter Phys 81: 061917

Uda S, Saito TH, Kudo T, Kokaji T, Tsuchiya T, Kubota H, Komori Y, Ozaki Y-I & Kuroda S (2013) Robustness and compensation of information transmission of signaling pathways. Science 341: 558–561

Wang G, Moffitt JR & Zhuang X (2018) Multiplexed imaging of high-density libraries of RNAs with MERFISH and expansion microscopy. Sci Rep 8: 4847

Welch JD, Kozareva V, Ferreira A, Vanderburg C, Martin C & Macosko EZ (2019) Single-Cell Multi-omic Integration Compares and Contrasts Features of Brain Cell Identity. Cell 177: 1873–1887.e17

Welf ES, Danuser G. Using fluctuation analysis to establish causal relations between cellular events without experimental perturbation. Biophys J. 2014;107(11):2492–2498.

Wong VC, Mathew S, Ramji R, Gaudet S & Miller-Jensen K (2019) Fold-Change Detection of NF-κB at Target Genes with Different Transcript Outputs. Biophys J 116: 709–724

Yao J, Pilko A & Wollman R (2016) Distinct cellular states determine calcium signaling response. Mol Syst Biol 12: 894

Zhang M, Eichhorn SW, Zingg B, Yao Z, Cotter K, Zeng H, Dong H & Zhuang X (2021) Spatially resolved cell atlas of the mouse primary motor cortex by MERFISH. Nature 598: 137–143

Zhang Q, Gupta S, Schipper DL, Kowalczyk GJ, Mancini AE, Faeder JR & Lee REC (2017) NF-κB Dynamics Discriminate between TNF Doses in Single Cells. Cell Syst 5: 638–645.e5

Zuo C & Chen L (2021) Deep-joint-learning analysis model of single cell transcriptome and open chromatin accessibility data. Brief Bioinform 22

